# Anticancer synthetic arylsulfonamides with Wnt1-modulating activity

**DOI:** 10.1101/2025.01.11.632297

**Authors:** Mihira Gutti, Jennifer Luo, Vihaan Sharma, Emilia Lee, Arshia Desarkar, Rushika Raval, Augustus Soedarmono, Joy Zhu, Edward Njoo

## Abstract

Dysregulation of the Wnt1/β-catenin signaling pathway has been demonstrated to be a driving factor in the propagation of several human cancers. Previous studies have discovered methyl 3-{[(4-methylphenyl)sulfonyl]amino}benzoate (MSAB) as a selective inhibitor of the Wnt1/β-catenin signaling pathway, which putatively functions through direct engagement of β-catenin. To understand how changes to the identity and position of the methyl ester affect the *in vitro* potency of this compound in Wnt1-driven mammalian cell lines, we prepared and evaluated three analogs of MSAB with 3- and 4-substituted methyl and ethyl esters. In MTT assays, analogs with methyl esters showed significantly more activity than their ethyl ester counterparts and both 4-substituted esters exhibited significantly attenuated antiproliferative activity, with MSAB exhibiting dose-dependent activity across cancerous cell lines. Further analysis by flow cytometry reveals low-Annexin V signal, suggesting that these compounds do not function via a pro-apoptotic pathway. Additionally, through a TCF/LEF-activated luciferase reporter cell assay, we observe that the 4-substituted methyl ester analogous to MSAB exhibits slightly diminished Wnt1-inhibitory activity, while 3- and 4-substituted ethyl esters exhibit minimal Wnt1-inhibitory activity. This difference in potency with a simple ester substitution might be attributed to several factors that ultimately drive antiproliferative activity, prompting the investigation of other potential substituents to further investigate the structure-activity relationship of these compounds as Wnt1-based antiproliferative agents.

## Introduction

The Wnt1/β-catenin signaling pathway performs critical functions in embryonic development,^1–3^ organogenesis,^4–9^ tissue homeostasis,^10,11^ and cell survival,^12,13^ through carefully maintained dynamic homeostasis of β-catenin. Activation and nuclear translocation of β-catenin triggers TCF/LEF transcription factors, which directly modulate several downstream target genes that are associated with cell survival, proliferation, and differentiation, including c-Myc and cyclinD1.^14–17^ Endogenous activation of the membrane-bound Wnt1/Frizzled complex leads to activation of the protein Disheveled, which is the direct modulator of a GSK-3β-driven β-catenin destruction complex. Phosphorylative ubiquitinylation and the proteasomal destruction of β-catenin in normal cells mediated by this system prevents uncontrollable proliferation^18,19^ **(Figure 1a).** On the other hand, dysregulation of this equilibrium caused by aberrant β-catenin activation has been previously demonstrated to be a driving factor in the propagation of several human cancers,^14,20,21^ including lung cancer,^22^ breast cancer,^23^ colorectal cancer,^20,24^ and melanoma.^25,26^

**Figure 1:**
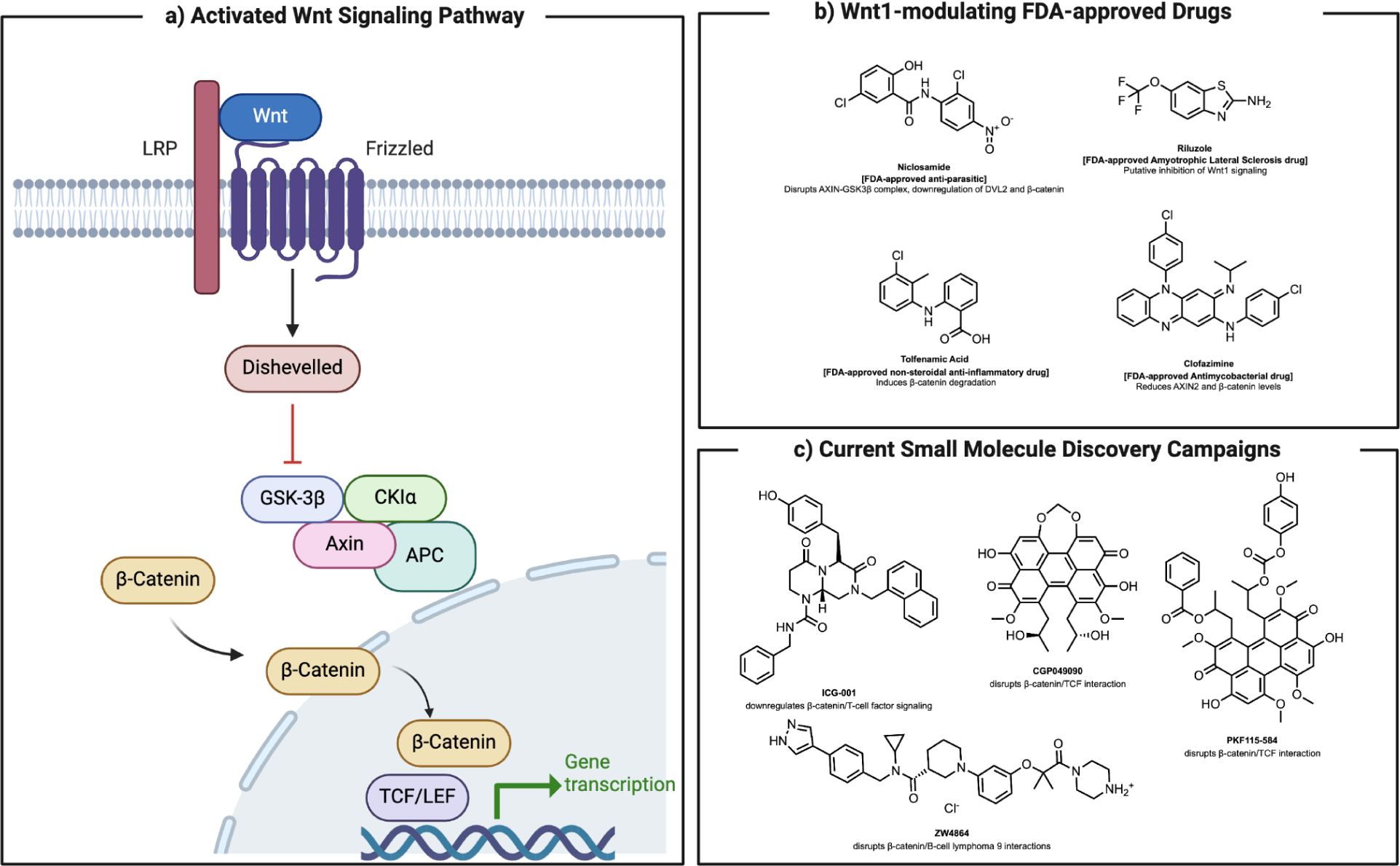
(a) Activation of the Wnt1/β-catenin signaling pathway inhibits GSK-3β, which causes nuclear translocation of β-catenin, triggering TCF/LEF transcription factors; (b) FDA-approved drugs with reported Wnt1/β-catenin modulatory activity include Niclosamide, Riluzole, Tolfenamic Acid, and Clofazimine; (c) Small molecules being developed as Wnt1/β-catenin modulators include ICG-001, ZW4864, PKF115-584 and CGP049090.

As such, ligands that modulate the Wnt1/β-catenin pathway have demonstrated strong preclinical promise as anti-cancer agents. Key examples of Wnt1-modulating FDA-approved drugs include Niclosamide, an anti-parasitic which has been reported to disrupt the AXIN-GSK-3β complex and downregulate DVL2 and β-catenin, and Riluzole, an Amyotrophic Lateral Sclerosis drug reported to inhibit Wnt/β-catenin/TCF-LEF (**Figure 1b)**.^27,28^ While several FDA-approved drugs and bioactive natural products exhibit partial Wnt-modulatory activity, no therapeutics selectively targeting Wnt1 signaling for the treatment of cancers are clinically available to date.^26,29^ To address this, several small molecule discovery campaigns have been initiated to target specific proteins along the Wnt1/β-catenin signal transduction relay: ICG-001,^30^ which binds to cyclic AMP response element-binding protein (CBP) and thus downregulates β-catenin/T-cell factor signaling; ZW4864,^31,32^ which disrupts β-catenin/B-cell lymphoma 9 interactions; and both PKF115-584 and CGP049090,^33,34^ which disrupt the β-catenin/TCF interaction (**Figure 1c)**.

Through a high-throughput small molecule screen, Hwang and coworkers previously discovered methyl 3-{[(4-methylphenyl)sulfonyl]amino}benzoate (MSAB **[1]**) as a selective inhibitor of the Wnt1/β-catenin signaling pathway. MSAB **[1]** putatively functions through direct engagement of β-catenin, stimulating its degradation and thus lowering its nuclear translocation.^35^ In a subsequent study, Magno and others have further elaborated on the structure-activity relationship of arylsulfonamide β-catenin inhibitors, showing that shifting the ester from the 4- to 2-position resulted in comparable β-catenin inhibitory activity.^36^ While the activity of MSAB **[1]** against Wnt/β-catenin-driven cancers is clear, more recent studies on the exact mechanism of small molecule β-catenin ligands, enabled by saturation transfer difference (STD) NMR spectroscopy experiments, have called into question the selectivity of several of these compounds in direct or indirect β-catenin modulation.^37^ This and previous studies further highlight the need for lead optimization of MSAB **[1]** and its analogs to determine the structural basis for both potency and target specificity.

To understand how changes to the identity and position of the methyl ester affect the *in vitro* potency of this compound in Wnt1-driven mammalian cell lines, we prepared and evaluated three analogs of MSAB **[1]** with 3- and 4-substituted methyl and ethyl esters **(Figure 2)**. We elected to prepare the corresponding ethyl ester given that there is relatively little known about the impact of the addition of a single carbon atom to the ester chain.^38^ Next, to interrogate the effect of repositioning the *meta-*methyl carboxylate ester to the *para* position, we prepared the corresponding 4-methyl carboxylate and 4-ethyl carboxylate esters. While these compounds have been previously synthesized,^39–41^ functional studies on these have been limited to chemical and physical properties, such as inhibiting metal corrosion,^41^ and relatively little is known about their biological activities. Through this study, we observed that substitution of the methyl ester to the ethyl ester, or changes to the position of the arylsulfonamide group generally results in a decrease in antiproliferative activity across several mammalian cell lines, demonstrating that structural changes of these types are not well-tolerated in Wnt-driven cell lines.

**Figure 2:**
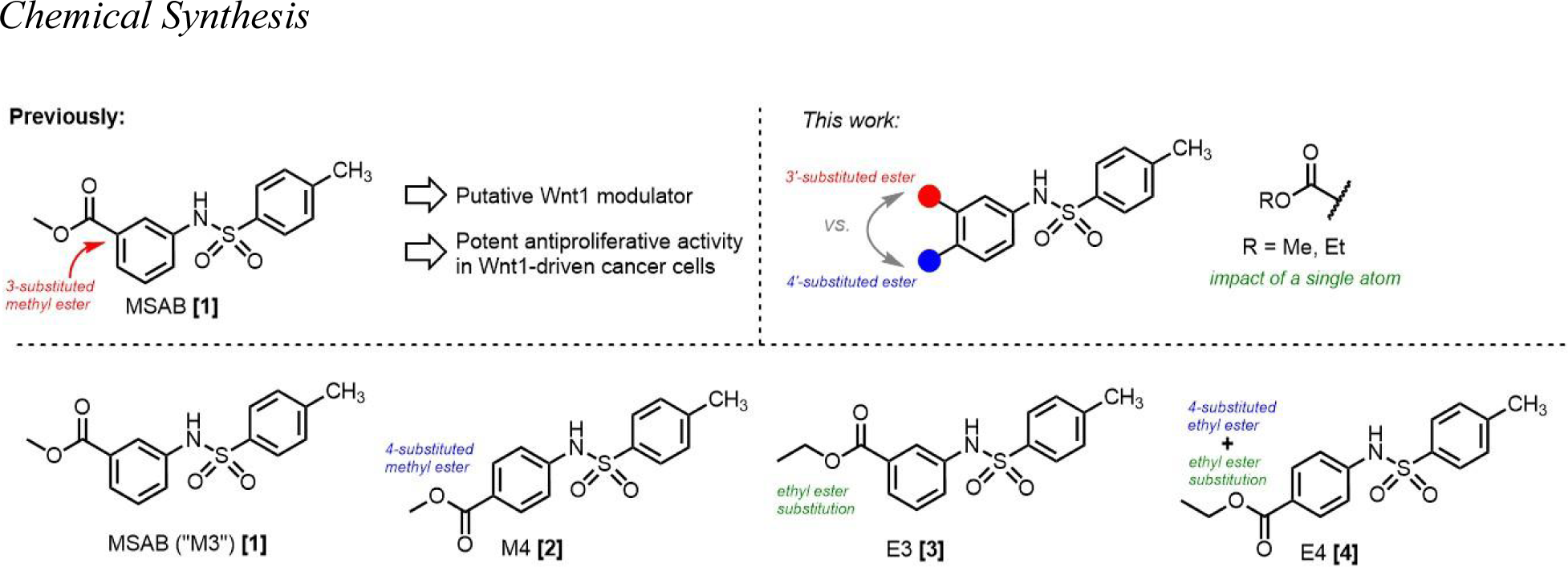
MSAB **[1]** and a series of three analogs with 3- and 4-methyl and ethyl ester substitutions.

In order to further assess the biological activity of MSAB **[1]** and its analogs, we conducted flow cytometry experiments to determine the potential involvement of pro-apoptotic mechanisms in cytotoxicity. Moreover, to examine whether the structure-activity relationship observed by cell viability assays is connected to inhibition of the Wnt1 signaling pathway, we employed a stable transfected reporter cell line that expresses firefly luciferase in the presence of TCF/LEF promoters. Through this, we determined that MSAB **[1]**, bearing a 3-methyl ester motif, exhibits GSK-3β independent inhibition of Wnt1 signaling. Collectively, these studies provide a blueprint for future analogs to further investigate the SAR and mechanism of these and similar arylsulfonamides as antiproliferative agents.

## Materials and Methods

### Materials

All reagents, solvents, and analytical standards were purchased from AK Scientific and Sigma Aldrich, and used without further purification. McCoy’s 5A Media, Dulbecco’s Modified Eagle’s Medium (DMEM), RPMI-1640 Medium, penicillin-streptomycin, and 0.25% trypsin EDTA were all obtained from Tribioscience (Sunnyvale, CA). Fetal bovine serum (FBS) was obtained from Gibco. Insulin and MTT reagents were obtained from AK Scientific.

### Cell Culture

Human colorectal cancer (HCT-116 and HT-29) cell lines were obtained from the European Collection of Authenticated Cell Cultures and maintained in McCoy’s 5A Media, supplemented with 10% fetal bovine serum (Gibco) and 1% penicillin-streptomycin (Tribioscience). The murine colorectal carcinoma (CT-26) cell line (ATCC CRL-2638) and maintained in RPMI-1640 Medium (Corning), supplemented with 10% fetal bovine serum and 1% penicillin-streptomycin. MDA-MB-231 human breast cancer cells were generously donated from SunnyBay Biotech (Fremont, CA) and cultured in RPMI-1640 (Corning), supplemented with 10% fetal bovine serum and 1% penicillin-streptomycin. Human embryonic kidney (HEK-293TN) cell lines were obtained from System Biosciences (Palo Alto, CA), and maintained in DMEM, supplemented with 10% fetal bovine serum, and 1% penicillin-streptomycin. Leading Light® Wnt Reporter 3T3 mouse embryonic fibroblast cells from Enzo Life Sciences (Cat. # ENZ-61002-0001) were maintained in DMEM, supplemented with 10% fetal bovine serum (FBS) and 1% penicillin-streptomycin. All cells were cultured in 25 cm^2^ and 75 cm^2^ flasks, in a humidified incubator at 37°C in 5% CO_2_.

### MTT Assay

Cells (HCT-116, HT-29, CT-26, MDA-MB-231P, HEK-293) were seeded in a 96-well tissue culture-treated flat bottom plate at 30% confluency and incubated at 37°C in 5% CO_2_. Cell viability was evaluated at 48 and 72 hours at different concentrations of compounds **[1]**-**[4]** in DMSO (500 µM, 250 µM, 50 µM, 25 µM, 5000 nM, 2500 nM, 500 nM, 250 nM). After incubation, 10 μL of a 5 mg/mL MTT solution in 1X PBS was added to each well, and incubated for an additional 2 hours at 37 °C. Then, the media was aspirated off, and the resulting formazan was solubilized with 100 μL of DMSO and incubated for 15 minutes before measuring absorbance at 570 nm using a Molecular Devices SpectraMax 250 Microplate Spectrophotometer. Percent cell viability was determined through the ratio of absorbance of treated cells to the negative control cells (DMSO, without compound). IC_50_ values were calculated on GraphPad Prism 10.

### Apoptosis Flow Cytometry Assay

Cells (HT-29, CT-26) were seeded at 70% confluency in a 12-well tissue culture-treated flat bottom plate and incubated in a humidified incubator at 37°C in 5% CO_2_. Treatment was conducted at 72 hours at the highest concentration of 500 µM. After incubation, the supernatant media was aspirated off, and the cells were trypsinized with 400 µL 0.25% EDTA-trypsin. Subsequently, the trypsin was deactivated with 400 µL of cell-line respective media (DMEM for HT-29, RPMI-1640 for CT26), and cells were pelleted by centrifugation. The cell pellet was then washed with 100 µL of 1X PBS, re-pelleted by centrifugation, and stained with 100 µL 1X Annexin binding buffer (Invitrogen), 5 µL FITC-Annexin stain (Southern Biotech), and 1 µL 1.0 mg/mL propidium iodide (PI) stain on ice for 15 minutes. Apoptosis analysis was performed by flow cytometry on a BD C6 Accuri Flow Cytometer to 25,000 events for each sample. Single cells were gated to exclude debris and cell aggregates based on forward scatter and side scatter. Based on Annexin V/PI quadrant gating, the percentage of cells that were viable, in the early apoptotic, late-stage apoptotic, and necrotic populations were quantified and expressed as percentages of total cell populations for each sample.

### Wnt1 Luciferase Reporter Cell Assay

Leading Light® Wnt Reporter 3T3 mouse embryonic fibroblast cells were obtained from Enzo LifeSciences (cat. ENZ-61002-0001) and seeded at 80% confluency in 96-well flat bottom plates (Corning Costar). After 24 hours of incubation at 37°C with 5% CO_2_, the plates were treated for 24 hours with **[1]-[4]** at various concentrations (500 µM, 250 µM, 50 µM, 25 µM) and CHIR-99021(AK Scientific) at 10 µM and 0 µM. A negative control of 0.5% v/v DMSO and a background control were used to establish background signal intensity in the absence of an inhibitor, and background signal in the absence of luciferin chemiluminescence. After 24 hours of incubation at 37°C with 5% CO_2_, 70 μL of Triton X-100 Lysis Buffer (0.1082 M Tris-HCl powder, 0.0419 M Tris-base powder, 75 mM NaCl, 3 mM MgCl_2_, 0.25% Triton X-100, H_2_O) was added to each well on top of the cell media. Then, 70 μL of the cell lysate was transferred to a black wall, flat bottom opaque 96-well plate (Corning Costar). 70 μL of the 3X Firefly Assay Buffer (15 mM Dithiothreitol, 0.6 mM coenzyme A, 0.45 mM ATP, 4.2 mg/mL D-luciferin, Triton X-100 Lysis Buffer) was added to the cell lysis and luminescence was quantified using a Tecan FarCyte Ultra Plate Reader.

## Results

### Chemical Synthesis

Compounds **[1]** to **[4]** were synthesized by sulfonylation of the corresponding aniline with *p-*toluenesulfonyl chloride according to the general procedure described: to an oven-dried round bottom flask fitted with a Teflon stir bar was added aniline **1a-1d** in the presence of 1-methylimidazole (3 eq.) and triethylamine (1 eq.) in dichloromethane (1.0 M). The crude reaction mixture was stirred at room temperature for 96 hours, and monitored by thin layer chromatography (TLC). The crude reaction mixture was worked up in a biphasic ethyl acetate/brine mixture, and the organic phases were collected and dried over anhydrous magnesium sulfate, filtered, and concentrated *in vacuo.* Subsequently, the crude residue was purified through silica gel flash column chromatography (0% → 40% ethyl acetate in hexanes). Compound identities and purities were established by ^1^H and ^13^C NMR spectroscopy and was consistent with previously reported characterization data.^35,39–41^

### Cell viability assays

We evaluated the antiproliferative activity of MSAB **[1]** and its analogs through MTT assays in human colorectal carcinoma (HCT-116), human colorectal adenocarcinoma (HT-29), murine colorectal carcinoma (CT-26), mammary triple-negative metastatic adenocarcinoma (MDA-MB-231), and human embryonic kidney (HEK-293) cell lines at 48 hour and 72 hour time points **(Figure 3)**. As expected, MSAB **[1]** exhibited dose- and time-dependent potency on cancer cell proliferation across all cell lines with IC_50_ values of 0.656, 1.20, 13.5, and 0.589 μΜ in HT-29, HCT-116, CT-26, and MDA-MB-231 cell lines at 48 hours, respectively **(Table 1)**. While MSAB **[1]** exhibited similar potencies in antiproliferative activity at 48 and 72 hour time points, compounds **[2]** and **[3]** were observed to generally bear greater antiproliferative activity at the 72 hour time point.

**Figure 3:**
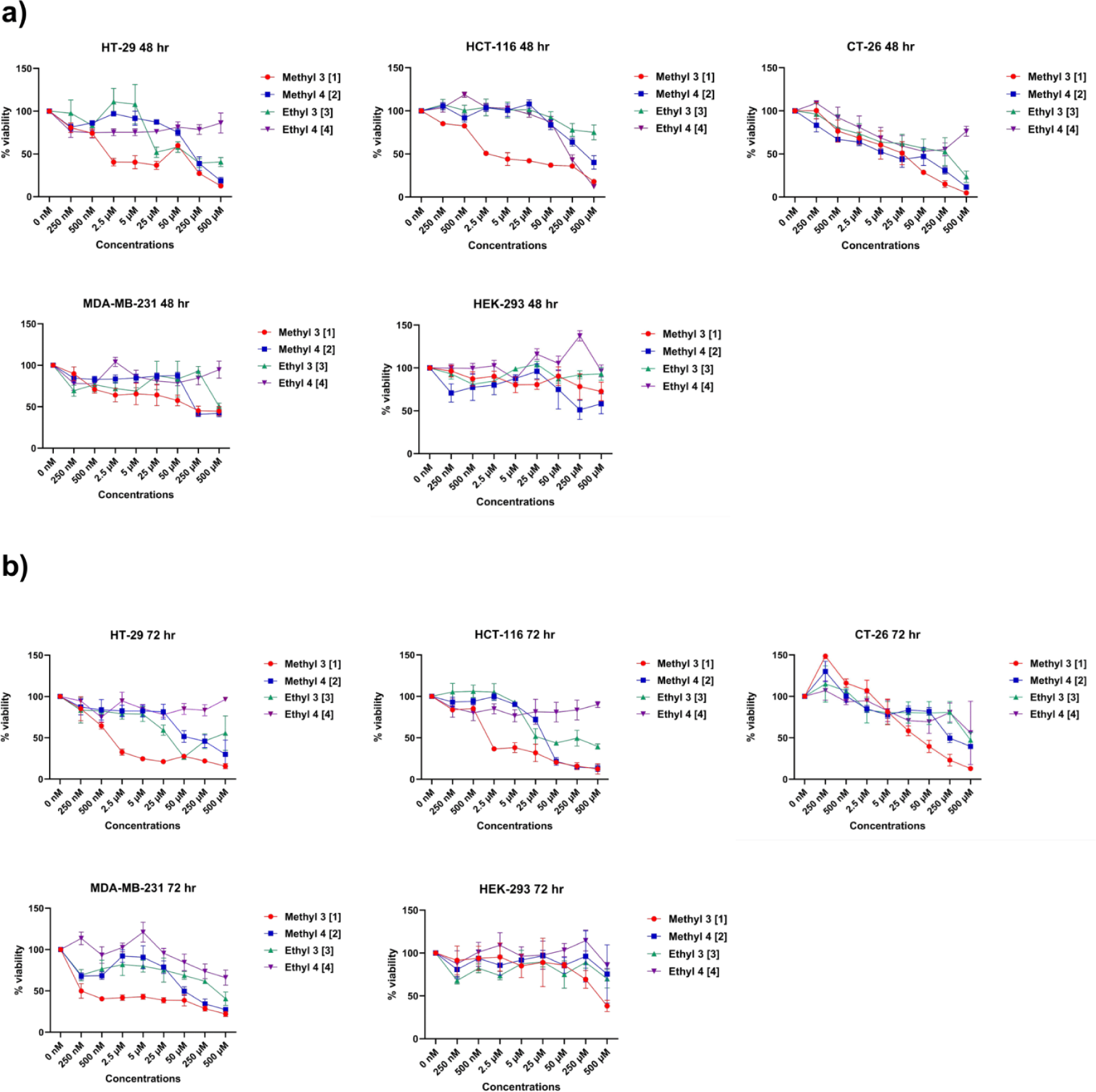
Cell viabilities across MSAB [1] and analogs [2]-[4] show that substitutions of the ester fragment are not tolerated. Cell viabilities of compounds **[1]-[4]** in HT-29, HCT-116, and CT-26 colon cancer cells, MDA-MB-231 breast cancer cells, and HEK-293 human embryonic kidney cells at **a)** 48 hour and **b)** 72 hour time course.

**Table 1:**
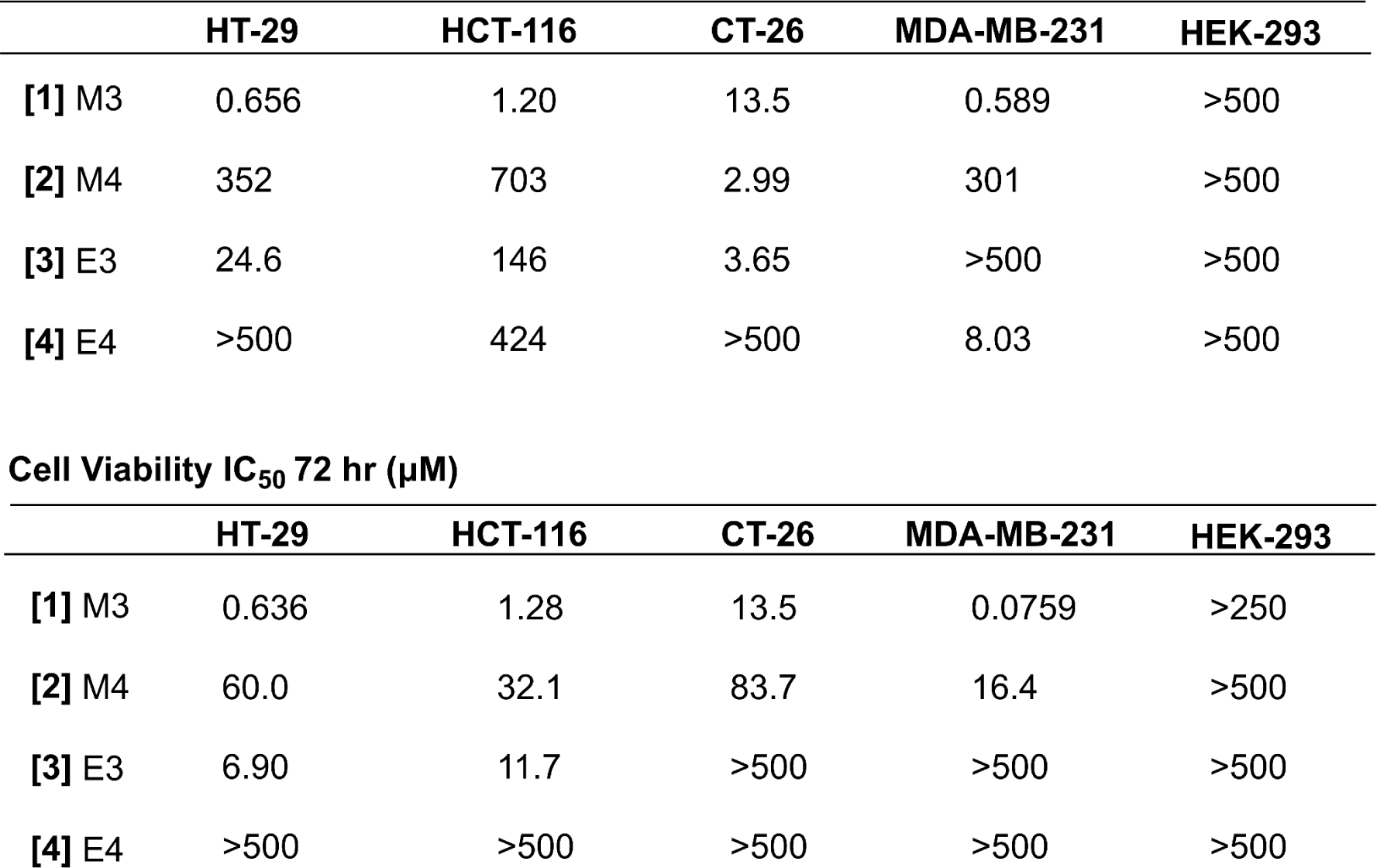
IC_50_ values of MSAB [1] and analogs [2]-[4]: MSAB **[1]** exhibited dose- and time-dependent potency on cancer cell proliferation across all cell lines. While MSAB **[1]** exhibited similar potencies in antiproliferative activity at 48 and 72 hour time points, compounds **[2]** and **[3]** were observed to generally bear greater antiproliferative activity at the 72 hour time point.

**Cell Viability IC<sub>50</sub> 48 hr (μM)**
|  | HT-29 | HCT-116 | CT-26 | MDA-MB-231 | HEK-293 |
| --- | --- | --- | --- | --- | --- |
| [1] M3 | 0.656 | 1.20 | 13.5 | 0.589 | >500 |
| [2] M4 | 352 | 703 | 2.99 | 301 | >500 |
| [3] E3 | 24.6 | 146 | 3.65 | >500 | >500 |
| [4] E4 | >500 | 424 | >500 | 8.03 | >500 |

**Table 1: IC<sub>50</sub> values of MSAB [1] and analogs [2]-[4]:** MSAB [1] exhibited dose- and time-dependent potency on cancer cell proliferation across all cell lines.
|  | HT-29 | HCT-116 | CT-26 | MDA-MB-231 | HEK-293 |
| --- | --- | --- | --- | --- | --- |
| [1] M3 | 0.636 | 1.28 | 13.5 | 0.0759 | >250 |
| [2] M4 | 60.0 | 32.1 | 83.7 | 16.4 | >500 |
| [3] E3 | 6.90 | 11.7 | >500 | >500 | >500 |
| [4] E4 | >500 | >500 | >500 | >500 | >500 |

The difference in potency in these cell lines indicates that the *in vitro* activity of MSAB **[1]** analogs is sensitive to the addition of a single carbon atom to the ester fragment or the repositioning of the ester fragment from the 3- to the 4-position.

### Flow Cytometry

We further sought to evaluate potential pro-apoptotic activity induced by our compounds through flow cytometry analysis. A standard Annexin V/Propidium Iodide protocol was followed to determine the percentage of cells undergoing apoptosis in cells treated with 500 μM of compounds **[1]-[4]** over a 24 and 72 hour period **(Figure 4)**. A common trend across all cell lines was elevated necrotic cells (PI stain), with MSAB **[1]** exhibiting the highest potency across all cell lines. The difference in necrotic cells from the control (24.13% for CT26, 8.44% for MDA-MB-231, 11.7% for HCT-116) demonstrates the antiproliferative properties of MSAB **[1]**. In CT-26 and HCT-116 cell lines, data was consistent with previous MTT cell viability assays, indicating that the *in vitro* activity of MSAB **[1]** analogs is sensitive to the repositioning of the ester fragment from the 3-position to the 4-position. However, the low Annexin-V signal across all the compounds suggests that there is limited involvement of pro-apoptotic mechanisms in cell death.

**Figure 4:**
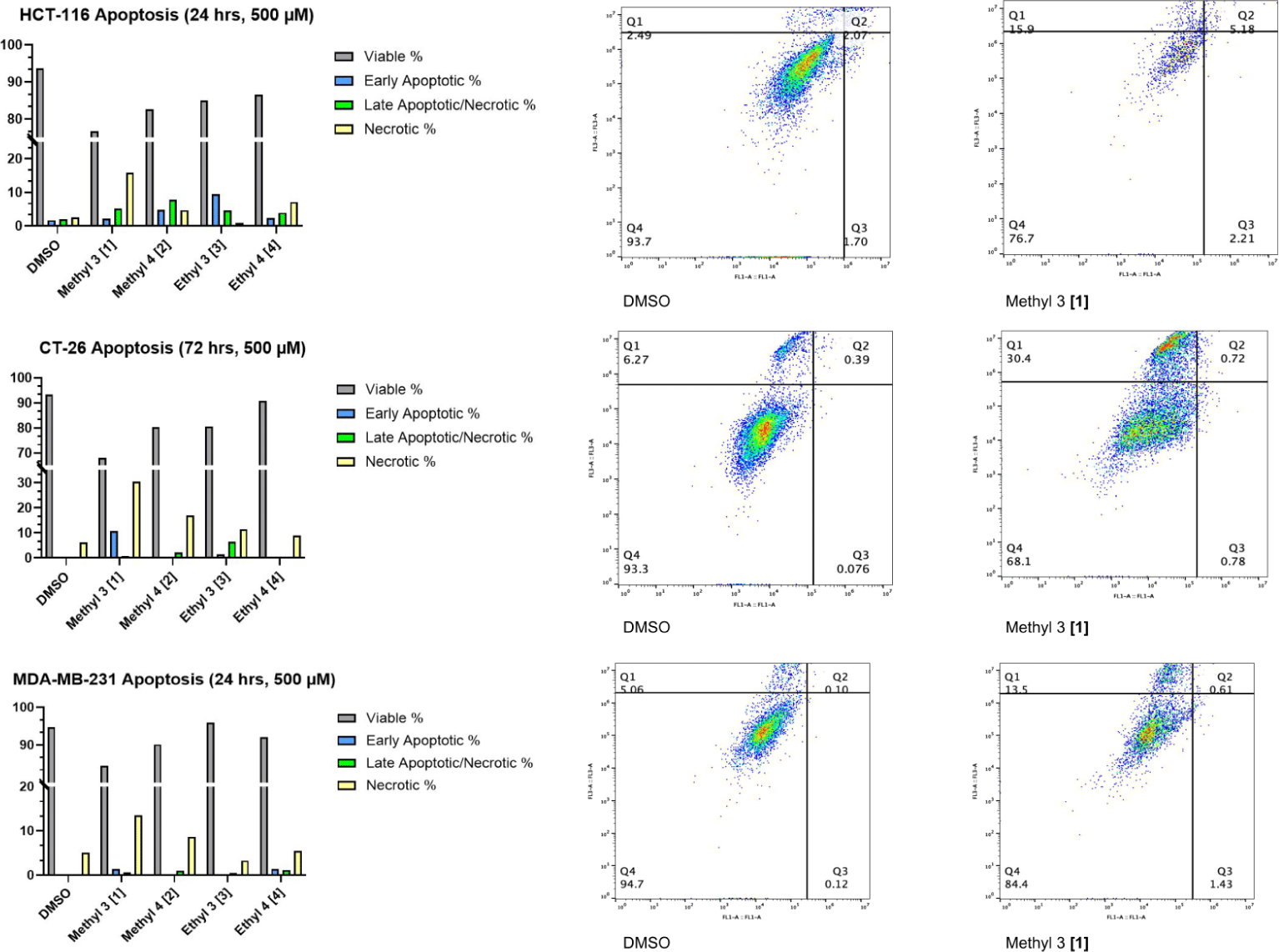
Flow cytometry of HCT-116, CT-26, and MDA-MB-231 cells treated with MSAB [1] and analogs. Elevated necrotic cells in compounds treated with MSAB **[1]** suggest that changes to the ester fragment are not well-tolerated. Standard Annexin-V/Propidium Iodide protocol was followed to determine the percentage of cells undergoing apoptosis in cells treated with compounds **[1]**-**[4]** and a DMSO negative control over a 24 and 72 hour period.

### Wnt1 Luciferase Reporter Cell Assay

Next, to determine the effect of compounds **[1]-[4]** on Wnt1 signaling, we employed an *in vitro* reporter cell luciferase assay using a stable transfected Wnt1-responsive 3T3 murine fibroblast cell line expressing a TCF/LEF-activated firefly luciferase gene. Compounds **[1]-[4]** were administered at 500 µM, 250 µM, 50 µM, 25 µM concentrations and incubated for 18 hours, after which a premade lysis buffer and luciferin substrate mix was added, and luminescence was observed on a 96-well plate reader. Consistent with their putative mechanism of action as Wnt1-pathway modulators, **[1]** and **[2]** exhibited potent inhibition of TCF/LEF-induced luciferase expression above 25 µM, whereas **[3]** exhibited inhibition above 50 µM. Compound **[4]** exhibited minimal activity in this assay, consistent with the low potency observed for this analog in cell viability experiments **(Figure 5a)**. Next, to evaluate whether this trend would hold upon inhibition of GSK-3β, we administered 10 µM CHIR-99021, a previously reported potent selective GSK-3β inhibitor,^42^ and found that while the baseline chemiluminescent signal was amplified as might be expected from GSK-3β inhibition induced amplification of TCF/LEF promoters, the same trends in dose-dependent decreasing Wnt1 activity was observed **(Figure 5b)**. Collectively, these results affirm the Wnt1-related mechanism of action in MSAB **[1]** and related aryl sulfonamides proposed by prior studies.^35,36^

**Figure 5:**
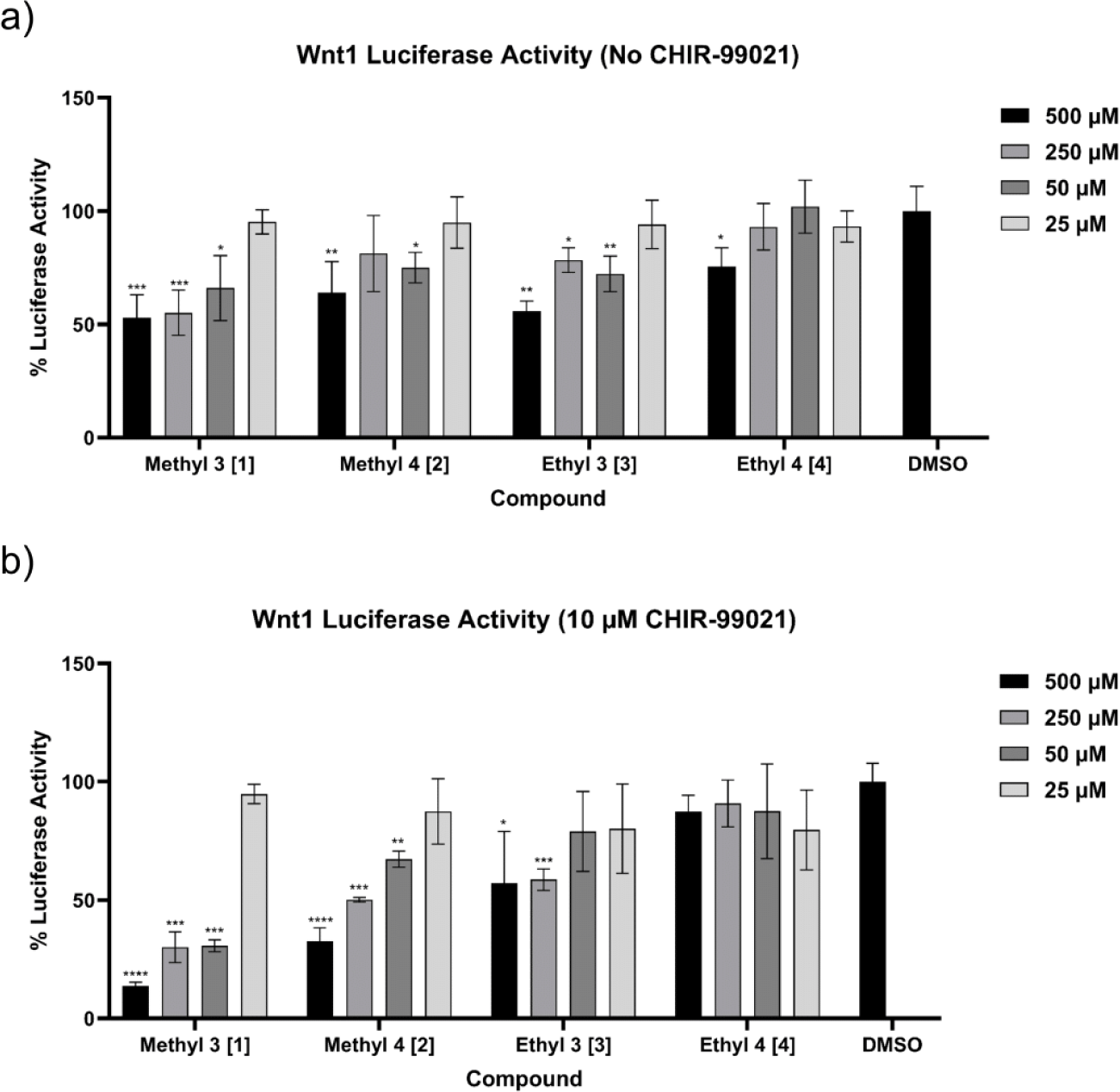
Activity of MSAB [1] and its analogs in a Wnt1-dependent luciferase reporter cell assay. Luciferase reporter activity from Leading Light® Wnt Reporter 3T3 mouse fibroblast cells was quantified through readout of the resulting luminescence from the oxidation of the luciferin substrate. A negative control was established with 0.5% v/v DMSO, and data is represented as means relative to a background control ± SD compared with the negative control using a Welch’s t-test (n = 4) (*P<0.05, **P<0.01, ***P<0.001, ****P<0.0001).

## Discussion

Motivated by the potency and selectivity of MSAB **[1]** in antiproliferative activity against Wnt1/β-catenin-driven cancers, we prepared and evaluated three analogs of **[1]**, including an ethyl ester variant and the corresponding analogs, where the 3-ester functionality is repositioned to the 4-position. As expected, we observed dose-dependent antiproliferative activity of MSAB **[1]** in HCT-116, HT-29, CT-26, and MDA-MB-231 cell lines with attenuated activity observed in non-cancerous HEK-293 cells. The corresponding ethyl ester **[3]**, while maintaining limited antiproliferative activity in HT-29, HCT-116, MDA-MB-231, and HEK-293 exhibited moderate loss in potency, indicating that increasing the alkyl-substituent may not be tolerated. The analogs with methyl esters showed higher potencies than their ethyl ester counterparts in all cell lines evaluated. Moreover, both 4-substituted methyl and ethyl carboxylate esters exhibited significantly reduced antiproliferative activity, suggesting that repositioning of the ester fragment is, too, not tolerated. While our analogs also exhibited dose-dependent antiproliferative activity to varying extents, MSAB **[1],** bearing the initial 3-substituted methyl ester originally reported by Hwang and co-workers, exhibited the most potent activity, while the analogous 4-substituted methyl ester **[2]** was observed to be slightly less potent. Additionally, we observed that simple substitution of the 3-methyl ester to the corresponding ethyl ester **[3]** was deleterious to antiproliferative activity in all cell lines, indicating that larger alkyl esters at this position may not be well tolerated. Compound **[4]** which bears both a 4-substituted ester fragment and the longer ethyl ester chain was observed to be the least potent among the set of compounds evaluated.

In order to determine potential apoptotic and necrotic mechanisms of cell death and antiproliferative activity, we analyzed the surface depolarization of phosphatidylserine, a marker for apoptotic activity, as well as membrane permeability to the DNA binding fluorophore propidium iodide (PI) via flow cytometry. In this assay, a similar trend of compound activity was identified; consistent with cell viability experiments, the 3-substituted methyl ester (MSAB **[1]**) provided potent and rapid increase in PI-positive cells.

Finally, to ascertain whether the biological activities of MSAB **[1]** and analogs **[2]** through **[4]** were connected to modulation of Wnt1 signaling as has been previously described, we employed a TCF/LEF-dependent luciferase reporter cell line, whose luminescence can be used as a quantitative proxy for Wnt1 pathway activation. Consistent to the patterns observed by initial cell viability assays and flow cytometry experiments, and consistent with prior reports, we observed a significant and dose-dependent decrease in Wnt1-driven luciferase signal when treated with MSAB **[1]**, and a proportionally lower Wnt1 inhibitory activity in 4-methyl ester **[2].** Expectedly, codelivery with 10 μM of the selective GSK-3β inhibitor CHIR-99021 led to higher overall signal intensities in this assay.

This difference in potency with a simple ester substitution, might be attributed to several other confounding factors in ultimate antiproliferative activity, including bioavailability, metabolic stability, and, nonspecific, off-target binding. Collectively, these results indicate that the most potent antiproliferative activity of MSAB **[1]** aryl sulfonamide analogs is found with 3-methyl carboxylate ester substituents and that even subtle structural changes to the alkyl ester substitution and positioning on aryl ester ring fragment are highly consequential in observed *in vitro* potency. This prompts the investigation of other potential substituents, including the design of analogs bearing modifications of the phenyl sulfonamide ring to further investigate the SAR of these compounds as Wnt1-based antiproliferative agents.

## Author Contributions

V.S., A.S., and J.Z. synthesized the compounds reported. M.G., J.L., E.L., A.D., and E.N. performed cell viability experiments and flow cytometry experiments. R.R. and E.N. performed the Wnt1 reporter cell assay. All authors contributed to writing of the manuscript.

## Acknowledgements

The authors express their gratitude to SunnyBay Biotech for supplying the HEK-293 and MDA-MB-231 cells and BJ Biosciences for supplying the CT-26 cells.

